# Fibromodulin Ablation Exacerbates the Severity of Acute DSS Colitis

**DOI:** 10.1101/2022.01.12.475640

**Authors:** Mariana Halasi, Mor Grinstein, Avner Adini, Irit Adini

## Abstract

Epidemiological studies have associated pigment production with protection against certain human diseases. In contrast to African Americans, European descendants are more likely to suffer from angiogenesis-dependent and inflammatory diseases, such as wet age-related macular degeneration (ARMD) and ulcerative colitis (UC), respectively. In this study, albino mice that produced high levels of fibromodulin (FMOD) developed less severe acute colitis compared with mice lacking in FMOD as assessed by clinical symptoms and histopathological changes. In a mouse model of dextran sodium sulfate (DSS)-induced acute colitis, FMOD depletion affected the expression and localization of tight junction proteins, contributing to the destruction of the epithelial barrier. Furthermore, this study revealed a stronger inflammatory response after DSS treatment in the absence of FMOD, where FMOD depletion led to an increase in activated T cells, plasmacytoid dendritic cells (pDCs), and type I IFN production. These findings point to FMOD as a potential biomarker of disease severity in UC among light-skinned individuals of European descent.

## INTRODUCTION

Ulcerative colitis (UC) is an inflammatory bowel disease (IBD) that affects primarily the mucosal surface of the large intestine (colon and rectum). It is characterized by destruction of the intestinal barrier, disappearance of the intestinal crypts, and infiltration of inflammatory cells ^1,2^. Epidemiological studies indicate the prevalence of UC continues to increase worldwide, particularly in developed countries. Individuals with North American and Northern European ancestry have the highest incidence and prevalence rates with 20/100,000 and 156 – 291/100,000 cases per year, respectively. Multiple factors contribute to the pathogenesis of UC, including environmental triggers, genetic predisposition, changes in intestinal microbiota, and immune system dysfunction ^3,4^.

UC progression is mainly affected by the dysregulation of tight and adherens junction proteins, which form the apical junction complex and maintain the epithelial barrier. Damage to these proteins results in a dysfunctional mucosal barrier and increased intestinal permeability ^5^. The molecular composition of the tight junctions (TJs) includes transmembrane proteins (claudin, occludin), cytoplasmic proteins (zonula occludens-1, ZO-1), and cytoskeletal components ^6^. The tight junctions serve to prevent the internalization of intestinal pathogens. Interestingly, dendritic cells (DCs), the most effective antigen-presenting cells, not only play a key role in maintaining immunological homeostasis by orchestrating innate and adaptive immune responses, but also express TJs. DCs are found in the intestinal mucosa at the interface with the external environment, where they act as sentinels for external pathogens ^7-9^.

Fibromodulin (FMOD), a small leucine-rich proteoglycan, plays a role in extracellular matrix (ECM) composition. FMOD was shown to activate the classical complement cascade by direct binding to C1q and by the deposition of C3b, C4b, and C9. FMOD can also silence the complement cascade by binding to factor H ^10,11^. Our previous work showed that FMOD up-regulates TGF-β1 secretion ^12-14^. TGF-β1 appears to enhance intestinal epithelial barrier function by inducing the production of TJ proteins, such as claudin-1, occludin, and ZO-1 ^15^. Furthermore, TGF-β1 is well known for its role in maintaining DCs in an immature state ^16^. During activation and migration of immature DCs to inflammatory sites, the DCs interact directly with ECM microenvironment proteins, such as FMOD, resulting in DC differentiation and maturation ^17,18^.

To advance our understanding of the pathogenesis of UC, we investigated the role of FMOD in a DSS-induced murine model of acute colitis, as well as in biopsy samples from healthy and diseased individuals with IBD.

## METHODS

### Mice

Mutant B6(Cg)-*Tyr^c-2J^*/J (FMOD+/+) 8-10-week-old male mice were purchased from the Jackson Laboratory. The mutant B6(Cg)-*Tyr^c-2J^*/J transgenic FMOD knockout (FMOD−/−) animals were established previously as described ^1^. The knockout animals were bred and maintained in pathogen-free conditions in the animal facility of Massachusetts General Hospital, Boston, MA. All animals were housed on 12 hr light-dark cycles under controlled temperatures, and they had free access to standard diet and water. All animal experiments were approved by and were conducted in accordance with the guidelines of the Institutional Animal Care and Use Committee at Massachusetts General Hospital.

### Induction of experimental DSS colitis

Acute colitis was induced in age-, and gender- matched, parental FMOD+/+ and FMOD−/− mice by the administration of 3% DSS (molecular weight 36,000-50,000; MP Biomedicals) ad libitum in autoclaved drinking water for 7 days. Body weight, stool consistency and bleeding were recorded daily to monitor colitis development. On day 7 after the start of DSS treatment, the animals were sacrificed, the gross pictures of the colon from cecum to rectum were taken, colon length was measured, and the colon tissue samples were processed for further downstream analysis.

### Human samples

Colon biopsy sections of healthy individuals and patients diagnosed with ulcerative colitis or Crohn’s disease were purchased from the Biorepository and Tissue Bank of the University of Massachusetts, Center for Clinical and Translational Science (UMCCTS).

### Scoring of bleeding

Bleeding score criteria follows as: 0= no blood; 1= small blood spot around anus; 2= visible rectal bleeding; 3= dried, chunks of blood on fur around anus.

### Scoring of Disease Activity Index (DAI)

DAI was calculated by the addition of the average weight loss score, the average stool consistency score and the average bleeding score and then divided by 3. Weight loss score criteria follows as: 0= no weight loss; 1= 1-5% weight loss; 2= 6-10% weight loss; 3= 11-20% weight loss. Stool consistency score follows as: 0= normal; 1= loose stool; 2= elongated loose stool; 3= diarrhea.

### Scoring of H&E sections

On the day of termination, the colon was removed, flushed with ice cold phosphate buffered saline (PBS) to remove feces and blood, cut longitudinally and prepared as a swiss roll for fixation in 4% paraformaldehyde (PF). Paraffin embedded sections were stained with Hematoxylin and Eosin (H&E) and histological scoring was performed in a blinded fashion independently by two pathologists. The histological sections were examined for the integrity of tissue architecture (0-4), ulceration (0-4), and submucosal edema (0-4).

### Total RNA extraction and quantitative real-time PCR (qRT-PCR)

Colon tissue samples were processed by the IBI Isolate reagent (IBI Scientific) for total RNA extraction. The High-Capacity cDNA Reverse Transcription Kit (Applied Biosystems) was used to synthesize complementary DNA (cDNA). Quantitative real-time PCR reactions were run on the LightCycler 480 instrument (Roche) using the PrimeTime Gene Expression Master Mix (IDT DNA) and the following mouse primers: Claudin1-S, 5′-ACA GCA TGG TAT GGA AAC AGA-3′, Claudin1-AS, 5′-AGG AGC AGG AAA GTA GGA CA-3; IFNα/β-S, 5′-TTC CTC AGC CAA GTT GCT-3′, IFNα/β-AS, 5′-GTG CAG TGT CCT AGT CCA G-3′; Ocludin-S, 5′-CAC TAT GAA ACA GAC TAC ACG ACA-3′, Ocludin-AS, 5′-GTT GAT CTG AAG TGA TAG GTG GA-3′; TJP1-S, 5′-AAT GAA TAA TAT CAG CAC CAT GCC-3′, TJP1-AS, 5′-GCC ACT ACA GTA TGA CCA TCC-3′; 18S-S, 5′-GAG ACT CTG GCA TGC TAA CTA G-3′, 18S-AS, 5′-GGA CAT CTA AGG GCA TCA CAG-3′; GAPDH-S, 5′-AAT GGT GAA GGT CGG TGT G-3′, GAPDH-AS, 5′-GTG GAG TCA TAC TGG AAC ATG TAG-3′.]

### Cell isolation and Flow cytometry

Mesenteric lymph nodes (mLN) of DSS treated animals were isolated, passed through a 100-µm cell strainer and red blood cells were lysed with the RBC lysis buffer (Biolegend). To differentiate between live and dead cells, cells were stained for 15 min at room temperature with the Zombie UV Fixable Viability Kit (Biolegend) as per company recommendation. Cell surface staining was performed for 30 min at 4ºC with fluorochrome-conjugated monoclonal antibodies to mouse antigens purchased from Biolegend: CD45 (30-F11), CD11c (N418), PDCA-1 (927), CD3 (17A2), CD4 (GK1.5), CD25 (PC61). Following staining cells were washed and fixed in 4% PF.

For intracellular staining, cells first were stained for cell surface markers, washed, then fixed, permeabilized and stained for intracellular transcription factor FoxP3 (MF-14; Biolegend) for 30 min at RT. Samples were analyzed by the FACSAria cell sorter (BD Biosciences), and the acquired data were analyzed using FlowJo software (version 10.6.1).

### Microscopy and Immunofluorescence staining

Brightfield images of H&E-stained sections were captured by Nikon Eclipse Ti2 microscope. For immunofluorescence staining, paraffin embedded colon sections were rehydrated by incubating in a series of xylene, ethanol (100%, 95%, 70% and 50%) and phosphate-buffered saline (PBS) solutions. After blocking, primary antibodies of CD45 (Biolegend), claudin-1 (BiCell Scientific), FMOD (Abcam) occludin (BiCell Scientific) or ZO-1 (BiCell Scientific) were applied for overnight incubation. As for secondary antibodies, Alexa Fluor 594 (Invitrogen) and Alexa Fluor 647 (Invitrogen) were utilized. To visualize nuclei, slides were mounted with VECTASHIELD Antifade mounting medium with DAPI (Vector Laboratories). Fluorescence imaging was performed with Nikon Eclipse Ti2 microscope or Leica SP8 X confocal microscope. Acquired images were analyzed using the NIS Elements software (Nikon) or the LASX life science software (Leica). Mean Fluorescence Intensity (MFI) was calculated by ImageJ 8.

### Statistical analysis

Statistical analysis was performed with Graphpad as described in the figure legends. *P* values of <0.05 were considered to be statistically significant.

## RESULTS

### FMOD expression is affected by UC in human subjects and mice

The expression of FMOD was assessed by immunostaining the colon biopsy samples from healthy control individuals and patients diagnosed with UC or Crohn's disease (CD). As shown in Fig.1A (left panel), in healthy individuals, FMOD was mainly detected in the brush border on the luminal cell surface of the colon. In contrast, a significant increase in FMOD expression was observed in the crypts of colon biopsy samples from UC patients (Fig.1A, right panel).

**Figure 1.**
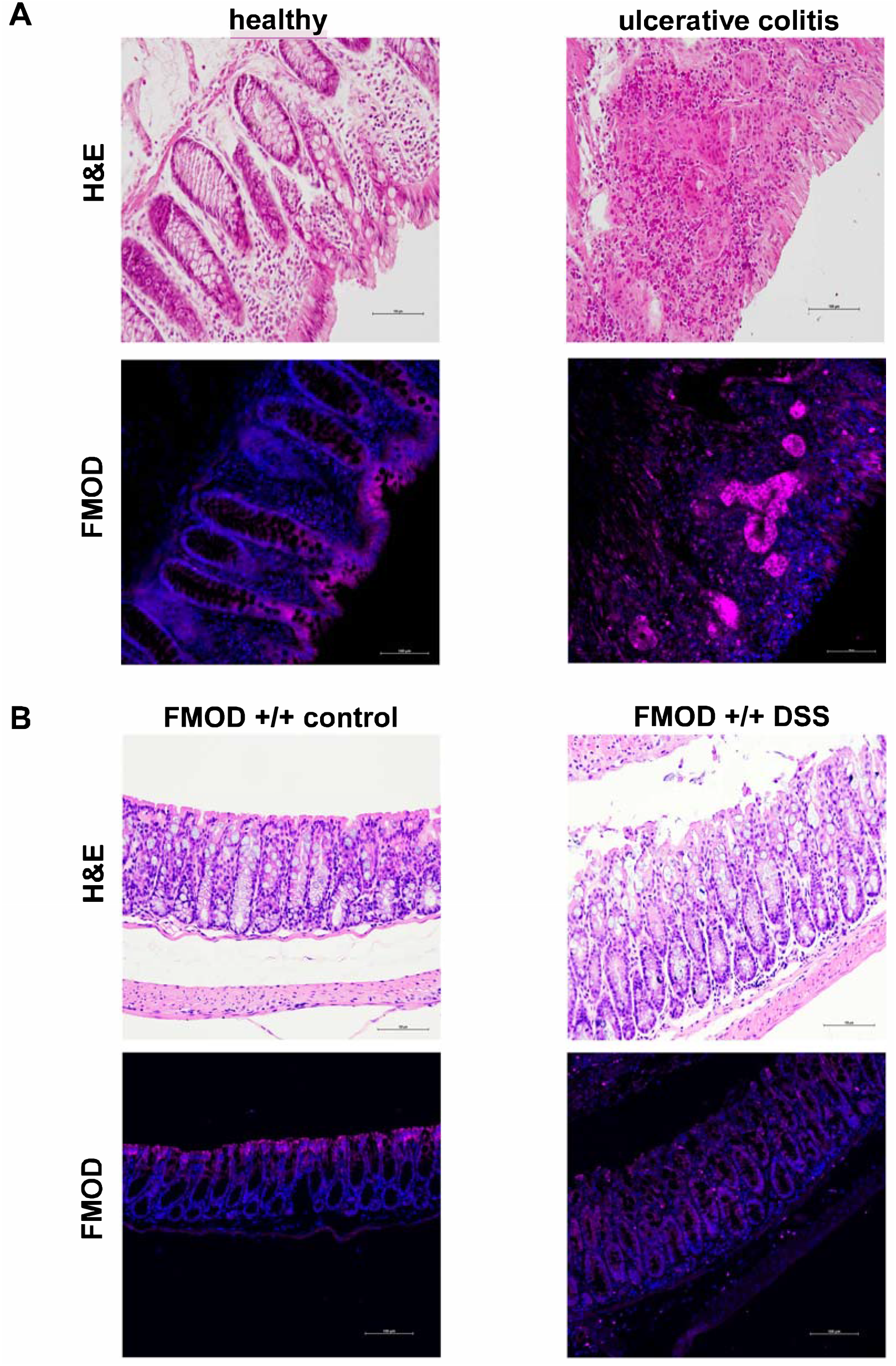
FMOD is upregulated and re-localized in colitis. **(A)** Colon biopsy samples from healthy individuals and patients diagnosed with UC were stained with H&E or subjected to immunostaining for FMOD. Representative images were captured at x20 magnification. **(B)** FMOD+/+ mice were given 3% DSS in drinking water for 7 days. Representative samples were H&E stained or immunostained for FMOD. Images were obtained at x20 magnification.

Since dextran sulfate sodium (DSS)-induced colitis is a well-established and commonly used animal model ^19,20^ that mirrors key aspects of human IBD, especially UC, FMOD level and localization was evaluated in animals with DSS-induced colitis. FMOD expression demonstrated the same distribution observed in the human samples (Fig.1B).

### Depletion of FMOD aggravated the DSS-induced clinical symptoms of colitis

To determine the role of FMOD in the development of acute colitis, age-matched transgenic FMOD knockout (FMOD−/−) male mice, along with FMOD+/+ animals, were exposed to 3% DSS for 7 days as described in Materials and Methods. The clinical signs of DSS-induced acute colitis, which include changes in body weight, bleeding, and fecal consistency, were recorded daily to monitor onset and progression of disease. Both FMOD+/+ and FMOD−/− DSS-treated experimental animals showed similar signs of acute colitis, including body weight loss (Fig. 2A) and shortening of the colon (Fig. 2B, C). However, DSS-treated FMOD−/− mice began losing weight 3 days after DSS was administered, while weight loss in FMOD+/+ animals began 5 days after DSS was administered. During the remainder of the DSS treatment, all of the treated animals showed substantial weight loss, amounting to 8.37% and 10% of their original weight in FMOD+/+ and FMOD−/− animals, respectively (Fig. 2A).

**Figure 2.**
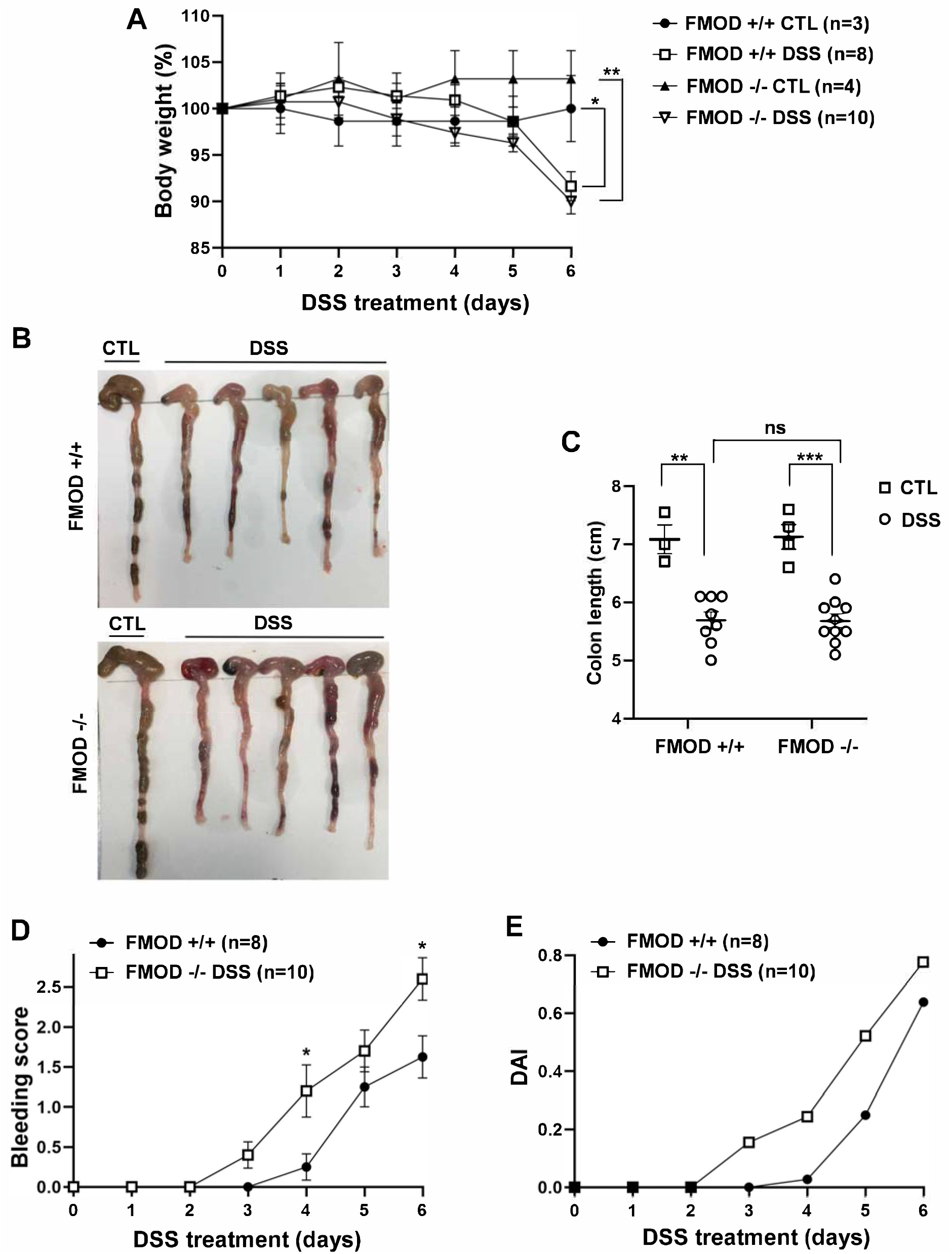
Ablation of FMOD facilitates DSS-related clinical signs of colitis. **(A-C)** FMOD+/+ (n=8) and transgenic FMOD−/− (n=10) mice were exposed to 3% DSS in drinking water for 7 days, and disease progression was monitored every day. Body weight was recorded daily **(A)**, gross pictures of the colons were taken **(B)**, and colon length was measured and plotted **(C)** on the day of sacrifice. **(D)** Rectal bleeding was recorded daily and scored according to the bleeding score criteria detailed in Materials and Methods after visually examining the anal area of treated animals. **(E)** DAI scores consisted of the sum of the average weight loss score, the average stool consistency score, and the average bleeding score divided by 3. Data are the mean◻±◻SEM from two independent experiments. Statistical analysis was performed by unpaired, two-tailed *t* test with Welch’s correction for (D) and one-way ANOVA with Tukey’s multiple comparisons test for (A and C): \**P*<0.05, \*\**P*<0.001, \*\*\**P*<0.0001.

One of the hallmarks of chemically induced colitis is colon shortening. As with the weight loss findings, colon shortening was similar between the treated groups on the day of necroscopy (19.71% in FMOD+/+ and 20.29% in FMOD−/−) but was significantly shorter compared to the respective untreated controls (Fig. 2B, C). Furthermore, both DSS-treated groups were visually checked for signs of rectal bleeding. Like weight loss, bleeding onset occurred earlier in the DSS-treated FMOD−/− animals, which started bleeding on day 3, compared to day 4 in the FMOD+/+ mice (Fig. 2D). Nevertheless, rectal bleeding remained more prominent in the FMOD−/− animals over the period of DSS treatment. The disease activity index (DAI) score, (i.e., the sum of the average weight loss score, the average stool consistency score, and the average bleeding score divided by 3), as shown in Fig 2E, indicated the FMOD−/− animals showed signs of colitis earlier and developed more severe disease compared with FMOD+/+ mice.

Together, these results demonstrated that the severity and onset of clinical symptoms of colitis, including weight loss, rectal bleeding, and diarrhea, were exacerbated in the FMOD knockout animals.

### FMOD-ablated mice exhibited severe pathological changes of the colon after DSS treatment

In addition to the phenotypical signs of colitis, histopathological changes in the colon were examined to further evaluate the possible role of FMOD in DSS-induced acute colitis. For histological H&E staining, the flushed and longitudinally cut colons were prepared using Swiss roll technique to permit a fuller assessment of the colonic pathology compared to the limited scope of transverse or longitudinal sections. Histological observations were carried out to assess the structural integrity of the intestinal epithelium (Fig. 3A). Colon sections from animals in both groups exhibited a fully intact epithelium with well-defined crypts, no edema, no leukocyte infiltration in the mucosa or sub-mucosa, and no ulceration or erosions (Fig. 3A). In contrast, the colon sections of DSS-treated animals showed different degrees of gut barrier compromise in the FMOD+/+ and FMOD−/− groups. Typical histological changes, including the presence of mononuclear inflammatory cells, disappearance of goblet cells, degradation of the epithelium, ulceration, edema, and leukocyte infiltration in the mucosa and sub-mucosa, were noted at different levels among the DSS-treated groups (Fig. 3A).

**Figure 3.**
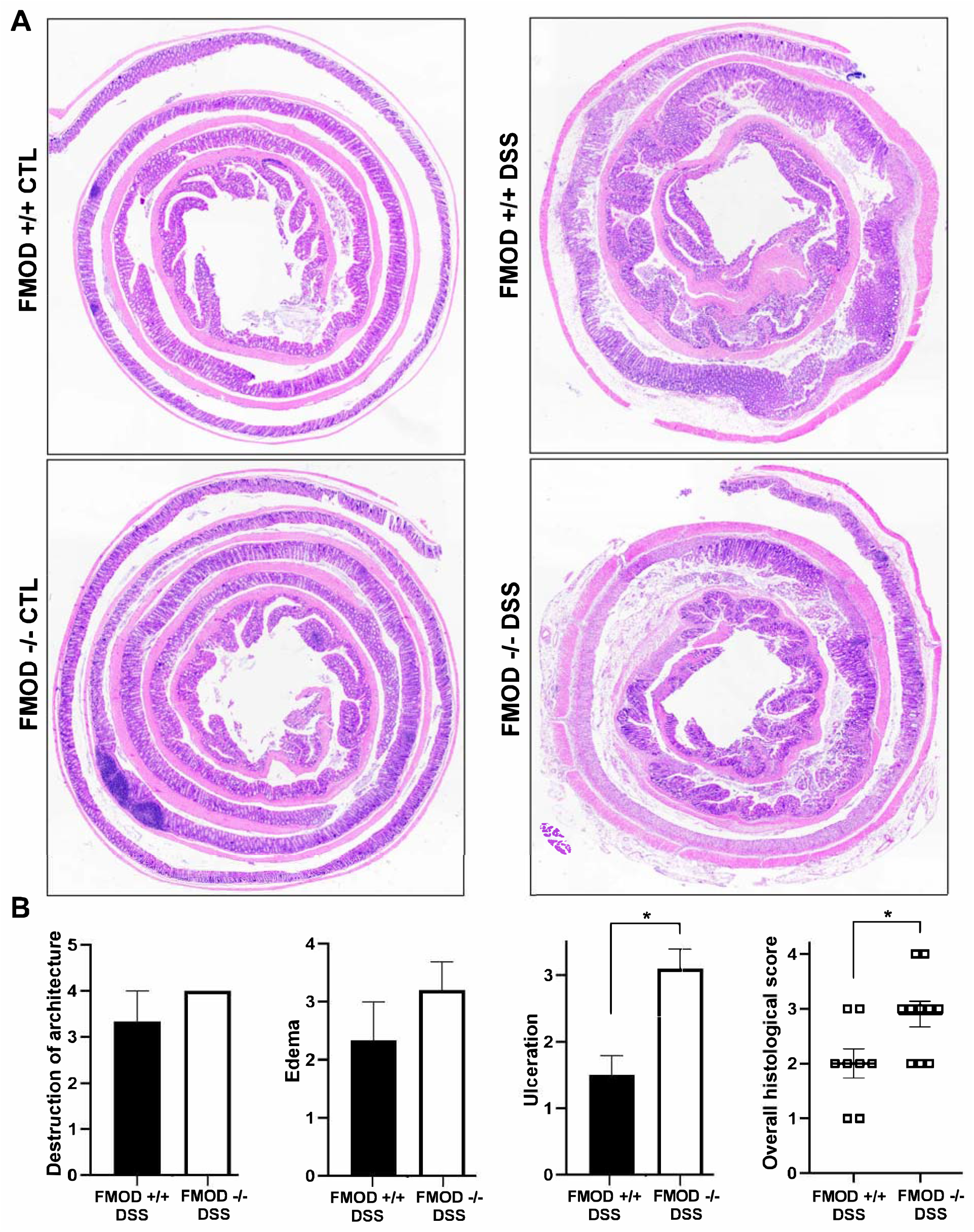
FMOD depletion facilitates DSS-induced pathological changes in the colon. **(A)** FMOD+/+ (n=8) and FMOD−/− (n=10) mice were given 3% DSS in drinking water for 7 days. Paraffin-embedded sections of the colon using Swiss roll technique were H&E stained. Images were obtained with x4 magnification. **(B)** Histological scores for the destruction of architecture (0-4), edema (0-4), ulceration (0-4), and overall score (0-3) of the full-length colon given in a blinded fashion are plotted. Data are represented as the mean◻±◻SEM of 3 control and 8 treated FMOD+/+, and 4 control and 10 treated FMOD−/− animals. \**P*<0.05 by unpaired, two-tailed *t* test with Welch’s correction.

In addition, the colon samples were scored blindly for the following criteria: destruction of architecture (0-4), ulceration (0-4), and edema (0-4) (Fig. 3B). The detailed quantitative histological analysis revealed more advanced disease in the DSS-treated FMOD−/− mice with an overall histological score average of 2.9 compared to the FMOD+/+ mice, which had an overall histological score average of 2 (Fig. 3B).

Taken together, these results suggested that although FMOD+/+ and FMOD −/− mice showed similar physical characteristics of acute colitis, they demonstrated marked differences in colonic histopathology after DSS treatment, in which the FMOD−/− animals showed higher susceptibility to DSS and developed more severe colitis. The histological analysis underscores FMOD’s role in moderating the damage inflicted by DSS on the epithelium, since knockout of FMOD resulted in substantial further damage.

### FMOD ablation facilitated epithelial barrier erosion

By its nature, DSS has the capacity to compromise the structural integrity of the epithelial cell barrier, possibly due to its interference with TJ proteins ^21^. To evaluate the effects of FMOD on the composition and expression of TJ proteins in DSS-induced colitis, colonic samples of untreated and DSS-treated groups were analyzed by immunofluorescence and qRT-PCR.

Our analysis focused on intracellular protein ZO-1 and transmembrane proteins occludin and claudin-1 on account of their well-established involvement in the impairment of intestinal barrier function in IBD disease models ^22,23^. ZO-1 protein expression was reduced, while claudin-1 expression was substantially upregulated (Fig. 4A) and occludin expression did not show a significant change (data not shown) in the DSS-treated colon sections of FMOD−/− mice compared to treated FMOD+/+ mice. Furthermore, quantification of fluorescence intensity by ImagJ 8 showed a 40% decrease in ZO-1 protein expression and a 1.6-fold increase of claudin-1 expression in DSS-treated FMOD−/− mice compared to treated FMOD+/+ mice (Fig.4B). To account for changes in protein expression levels, we also validated these findings at the mRNA level. The results showed that the mRNA level of ZO-1 was significantly reduced, claudin-1 mRNA was substantially upregulated, while occludin mRNA expression showed no statistically significant change in the DSS-treated FMOD−/− mice (Fig. 4C).

**Figure 4.**
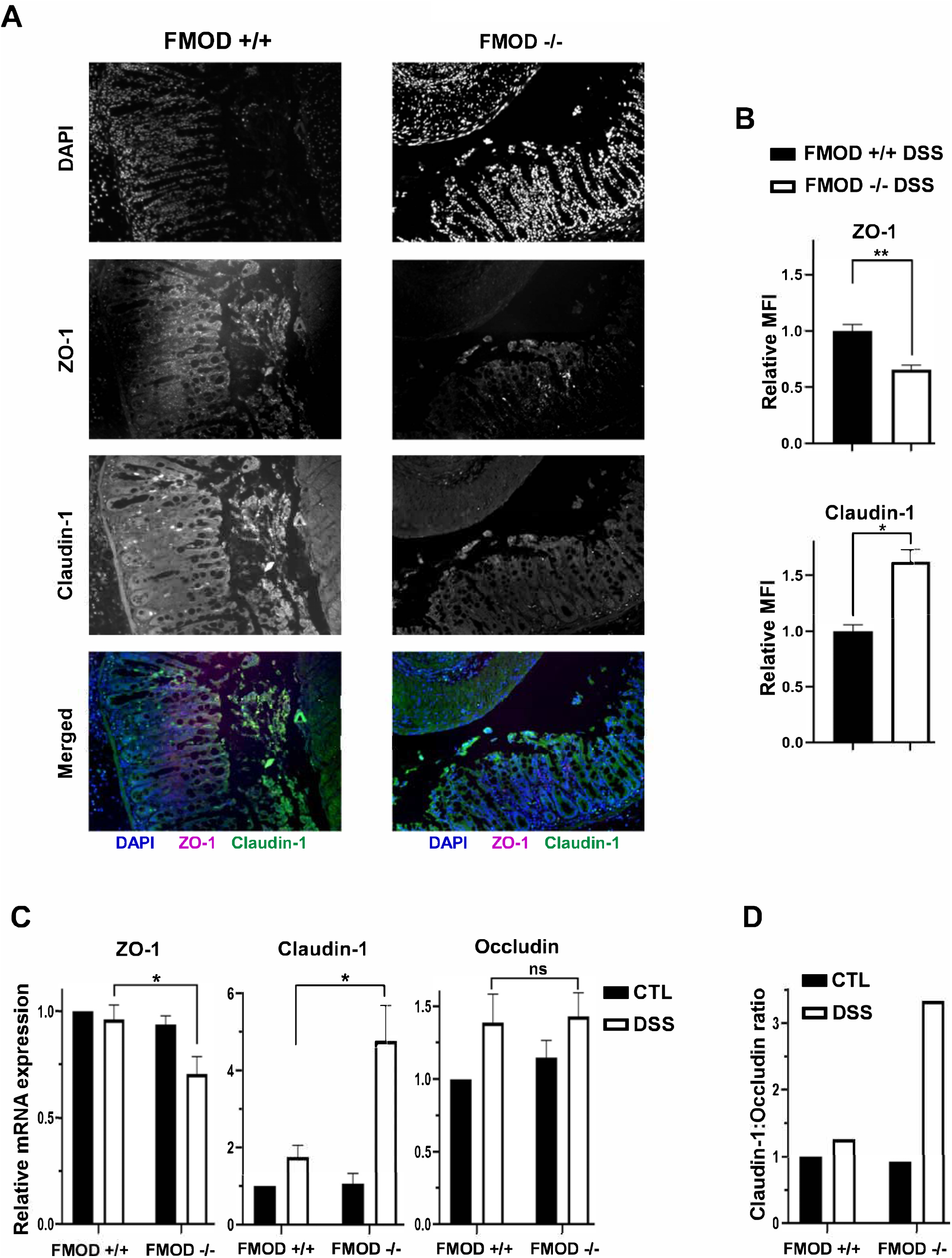
Knockout of FMOD amplifies the changes in expression of TJ proteins following DSS treatment. **(A)** Paraffin-embedded sections of DSS-treated colon Swiss rolls were stained with anti-mouse ZO-1 or claudin-1. Images were captured at x20 magnifications. **(B)** The graph shows the quantification of mean fluorescence intensity (MFI). **(C)** The transcriptional expression of TJ proteins ZO-1, claudin-1, and occludin was measured by qRT-PCR in the colon of untreated and DSS-treated FMOD+/+ and FMOD−/− mice. The graphs show quantification as fold of mRNA expression levels compared to untreated FMOD+/+ mice (n=3), mean◻±◻SEM 3 (FMOD+/+) or 4 (FMOD−/−) control and 8 (FMOD+/+) or 10 (FMOD−/−) treated mice. **(D)** The claudin-1 : occludin ratio was calculated using the respective mRNA expression levels. Statistical analysis was performed by unpaired, two-tailed *t* test with Welch’s correction: \**P*<0.05, \*\**P*<0.01

Poritz *et. al.^21^* found that occludin levels decrease and claudin-1 levels increase in the colon of human subjects with UC, resulting in a higher claudin-1: occludin (C:O) ratio compared to healthy colon. Using the C:O ratio calculated from the respective mRNA expression levels, we observed a 3-fold increase in disease severity in the absence of FMOD after DSS treatment (Fig. 4D).

### FMOD alleviated DSS-induced colonic inflammation

Leukocyte infiltration has been associated with the pathogenesis of human IBD. It has also been implicated in the acute phase of DSS-induced colitis ^24-26^. The presence of leukocytes in the colonic tissue was assessed by immunostaining for CD45 on colon sections from DSS-treated mice. Higher expression of CD45 (Fig. 5A) was observed in DSS-treated FMOD−/− animals, suggesting inflammation becomes more robust in the absence of FMOD.

**Figure 5.**
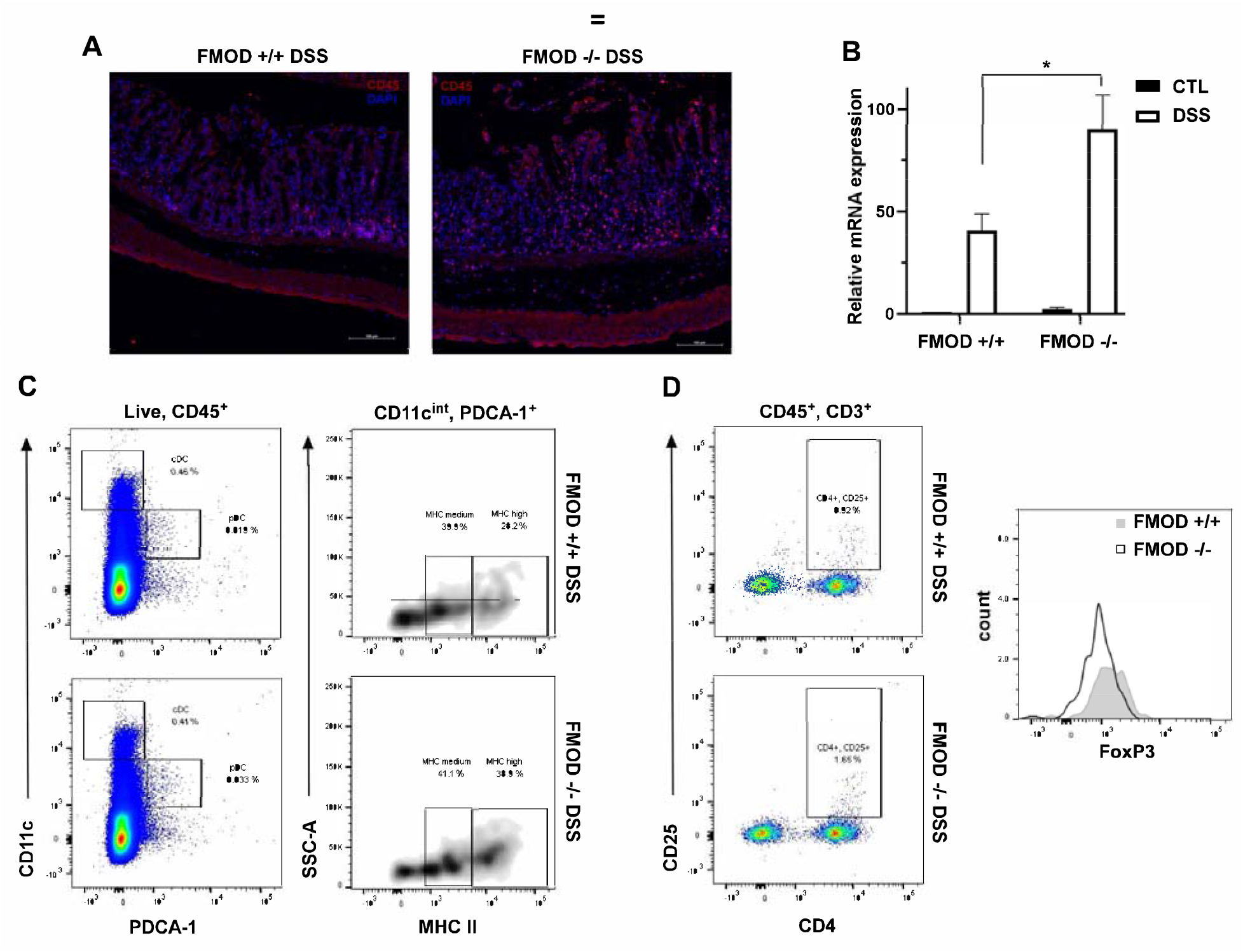
FMOD moderates DSS-induced colonic inflammation. **(A)** Paraffin-embedded sections of the DSS-treated colon Swiss rolls were stained with anti-mouse CD45. Images were captured at x20 magnifications. **(B)** The transcriptional expression of IFNα/β was measured by qRT-PCR in the colon. The graph shows quantification as fold of mRNA expression levels compared to untreated FMOD+/+ mice (n=3), mean◻±◻SEM 3 (FMOD+/+) or 4 (FMOD−/−) control and 8 (FMOD+/+) or 10 (FMOD−/−) treated mice. \**P*<0.05 by unpaired, two-tailed *t* test with Welch’s correction. **(C)** The mesenteric lymph nodes (mLN) of DSS-treated FMOD+/+ (n=5) and FMOD−/− (n=5) mice were analyzed by flow cytometry for the frequency (*left panel*) and the maturity (*right panel*) of pDCs. Data are shown as a dot plot (frequency) or a density plot (maturity), and numbers represent the percentage of CD11c^int^PDCA-1^+^ pDCs or CD11c^int^PDCA-1^+^MHC II^+^ pDCs among the live CD45^+^ leukocytes, respectively. **(D)** Cells from the mLNs of treated FMOD+/+ and FMOD−/− mice were stained for the expression of CD25 and FoxP3 and analyzed by flow cytometry. Dot plots show CD25 and CD4 staining pre-gated on live CD45^+^CD3^+^ cells. The histogram represents FoxP3 staining pre-gated on live CD45^+^CD3^+^CD4^+^CD25^+^ cells.

Colonic inflammation was further examined by assessing cytokine production in the colonic tissue and the immune response in mesenteric lymph nodes (mLNs), which are known to drain the colonic tissue^27^. First, the production of inflammatory mediators was measured in colon tissue samples by qRT-PCR. Markedly higher mRNA expression of immunomodulatory cytokine interferon α/β(IFN α/β) was observed in the DSS-treated FMOD−/− animals, underscoring the development of a potentially more robust inflammatory response (Fig. 5B).

To characterize the recruited and activated immune cell populations in systemic organs following DSS exposure, mLNs and spleens were harvested and analyzed by flow cytometry. A decrease in the splenic dendritic cell (DC) and T cell populations in both DSS-treated groups was detected (data not shown). Significantly more plasmacytoid dendritic cells (pDCs) with increased MHC II (major histocompatibility complex II) maturation markers on their surface were observed in the mLNs of treated FMOD−/− animals (Fig. 5C). Analysis of T cells revealed significantly more activated T cells in the DSS-treated FMOD−/− group, while increased mobilization of FoxP3+ T reg cells was observed in the FMOD+/+ DSS-treated mice (Fig. 5D), corroborating the presence of a stronger inflammatory response in the background of FMOD KO after DSS administration.

These results suggest that in the absence of FMOD, acute inflammation, represented by activated pDCs and reduced numbers of T regulatory cells, is significantly increased.

## DISCUSSION

Ulcerative colitis is an idiopathic IBD limited to the colon. It is a multifactorial in nature, characterized by dysregulation of the immune system, epithelial barrier defects, and changes in the intestinal microbiota. Based on our earlier findings ^12^, we hypothesize that FMOD, a prominent ECM protein, plays a role in inflammation and in the maintenance of the epithelial barrier. This study investigates the potential role of FMOD in the pathogenesis of UC utilizing the widely used DSS-induced acute colitis mouse model.

FMOD, a cytosolic protein with a known secretory sequence but no transmembrane domain, is characterized as a class II member of the small leucine-rich proteoglycan family ^28,29^. It was first described in human articular cartilage ^30^. Although it is ubiquitously expressed in all tissues among vertebrates, its expression is most abundant in the extracellular matrix (ECM) of the connective tissues, where it plays a key role in the organization of collagen fibrils ^31^. In recent years, additional tissue-specific activities have been reported, such as its role in muscle development ^32^, cell reprogramming ^33^, angiogenesis, and regulation of TGFβ-1 secretion ^12,34,35^. FMOD has also been associated with inflammatory diseases of the joint and anti-inflammatory responses in wound healing ^36^. To date, the role of FMOD in inflammatory and autoimmune diseases has been modestly explored. Increased expression of FMOD has been observed in patients with fibrotic kidney disease ^37^, heart failure, and atherosclerosis ^36^.

Through our experiments, we observed upregulated FMOD expression in both IBD patients and animals with DSS-induced acute colitis (Fig. 1). This observation may not be too surprising, since colitis causes oxidative stress, which is well-known to stimulate the expression of FMOD ^38^. Interestingly, however, deletion of FMOD in a transgenic acute colitis mouse model did not alleviate disease burden. Rather, it intensified the onset and severity of the clinical colitis symptoms as measured by weight loss, rectal bleeding, and DAI score (Fig. 2). In addition, histopathological examination of full-length colon sections underscored the development of more severe colitis in the absence of FMOD (Fig. 3).

The epithelial barrier physically separates the host immune cells in the intestinal mucosa from the luminal microbiota. Damage to the integrity of the colonic epithelia increases mucosal permeability due to the dysregulation of TJ proteins ^2,39^. Although, epithelial barrier erosion is a key feature of clinical IBD, it is still debated whether such dysfunction is a manifestation or a cause of UC ^40,41^. Loss and redistribution of tight junction proteins are commonly reported in acute DSS colitis ^21^. In agreement with these findings, marked differences in mRNA and protein expression of TJ proteins, ZO-1 and claudin-1, were observed between the DSS-treated groups (Fig. 4), showing more robust changes in the absence of FMOD.

Further investigation into the effect of FMOD depletion on colitis revealed the presence of a stronger inflammatory immune response in DSS-treated FMOD−/− mice (Fig. 5). Studies have shown that FMOD plays an important role in inflammatory responses through activating the classical and alternative complement pathways by directly binding to C1q and C3b ^10^. The activated complement system contributes to the adaptive and cellular immune responses by interacting with toll-like receptors (TLRs), antigen-presenting cells, and immune cells such as polymorphonuclear leukocytes, DCs, B cells, as well as T lymphocytes ^10,42^. Interestingly, the deposition of C1q can also inhibit molecular processes required for DC differentiation and IFN-α production by pDCs ^43-45^. Our results align with these previously described findings. Specifically, the presence of FMOD in FMOD+/+ animals, probably via complement activation, induced a more protective immune response as evidenced by the higher number of Foxp3+ T reg cells. Likewise, the absence of FMOD in FMOD−/− animals was associated with increased production of type I IFN, higher numbers of pDCs, and activated T cells indicating a stronger inflammatory response (Fig. 5).

Here we provide insight into the connection between FMOD and the increased susceptibility to colitis by employing genetic and biochemical approaches. In the acute phase of DSS-induced colitis, the presence of FMOD alleviates disease symptoms by maintaining the integrity of the epithelial barrier, reducing pDC activation, and inducing regulatory T cell responses. Future studies are warranted to examine the potential utility of tissue FMOD as a biomarker to gauge the severity of UC among light-skinned individuals of European descent.

## ACKNOWLEDGMENT

The authors thank Professors Rivka Dresner-Pollak, Martin L. Yarmush, MD, PhD and Yuval Tal, MD, PhD for fruitful discussions. This study was supported in part by a grant from NIH (RO1EY024046).

